# Ecological relationships between human gut bacteria predicted from analysis of dense microbiome time series data from US travelers in Bangladesh

**DOI:** 10.1101/2025.02.27.639550

**Authors:** Casey G. Martin, Laurie M. Lyon, Antonio Gonzalez, Rob Knight, Catherine Lozupone

## Abstract

Gut microbiomes provide critical host homeostatic functions, resulting from a complex web of ecological interactions among community members. We studied these interactions using a time-lagged correlational strategy of dense longitudinal sequence data from Western individuals traveling abroad to Bangladesh who experienced diarrhea. We identified both negative (140) and positive (78) relationships between bacterial pairs. Positive relationships occurred in pairs that were significantly more phylogenetically distant, such as inter-order associations between Clostridiales and Bacteroidales, while negative relationships were more between more phylogenetically related pairs. Further analysis of computationally predicted genome content and metabolic pathways revealed that cooperative bacterial pairs overlapped less in function and offered each other metabolic support, while competitive pairs were more likely to compete for the same resources. Predicted levels of B vitamins (B5 and B3), enoyl acyl-carrier protein (acp) reductase II (FabK) and its metabolites, and nucleotide/nucleoside derivatives were able to differentiate negatively and positively associated microbe pairs. Ultimately, our findings show that combining time-series analysis with metabolic/genomic network analysis can identify relationships between bacteria with plausible causal mechanisms that are consistent with existing ecological and biochemical observations.

**IMPORTANCE:** Understanding how microbes in the gut interact with each other is important for devising strategies to target the human gut microbiome therapeutically. For instance, understanding competitive relationships, where a shared need of similar limited resources limits the degree to which two microbes can co-exist, can inform strategies for limiting colonization of undesirable microbes. Understanding cooperative relationships, where one microbe provides the other with substrates needed for growth, can inform strategies to promote desirable microbes. By evaluating dense time-series gut microbiome data from individuals who experienced diarrhea while traveling, we were able to predict both cooperative and competitive relationships among human gut microbes as those whose abundances were significantly related within an individual over time. Strikingly, in subsequent analyses performed using inferred genomic information, pairs with negative associations from the time series analysis were predicted to compete over more metabolic substrates, and pairs with positive associations had significantly more metabolic complementarity. These predictions regarding the underlying molecular bases of interactions could inform how nutritional environment will impact interactions between gut microbiome community members.

## INTRODUCTION

Microbial communities play central ecological roles, ranging from environmental biogeochemical cycling to supplying integral homeostatic functions for their associated hosts across the animal, fungal, and plant kingdoms (1–3). In the human gut, ecosystem services provided by the microbiome include energy harvest, immune education, and pathogen exclusion, and are the result of a complex web of facilitative and antagonistic interactions between community members (4, 5). Even though the composition of the human gut microbiome has been extensively characterized across health and disease states, the fundamental ecological relationships between most consortium members have yet to be described. Resolving ecological linkages between microorganisms is important for the construction of basic microecological frameworks of community assembly and stability. Key questions include: How prevalent are cooperative and competitive relationships between bacteria in the gut and how do these interactions shape community composition? Are there phylogenetic patterns in these relationships? What attributes influence two microbes’ propensity to cooperate or compete? Questions regarding how ecological interactions like competition are influenced by evolutionary history and relatedness can be traced to back to Darwin, where he outlined his Congeneric Competition Hypothesis, a model that predicts that closely related organisms are more likely to share traits and occupy the same ecological niche, thereby leading to competition (6). Since the human gut has high functional redundancy where many microbes have highly overlapping metabolic pathways, competition due to niche overlap between functionally similar microbes may be an important force in shaping the composition and resilience of gut communities (7). On the other hand, metabolic cooperation has been documented in the gut, such as for utilization of host-derived glycoproteins or B-vitamins salvage (8, 9). Determining the importance and specific mechanisms of both competitive and cooperative relationships in the gut microbiome will be essential for devising strategies to promote healthy compositions.

While canonical experimental approaches are still the gold standard for in depth investigation of microbe-microbe interactions, high-throughput sequencing has enabled the advancement of statistical methods for inferring ecological interactions that can then be tested in the lab. For the past half-century, co-occurrence has been a popular tool in attempts to predict macroecological interactions, and these techniques have been widely adopted in high-throughput surveys of microbial relative abundance (10–12). The popularity of co-occurrence analysis is understandable, as it is amenable to a variety of experimental designs and cross-sectional data, but co-occurrence is not a reliable proxy of a direct interaction because it can occur between microbes with similar environmental preferences (13). Co-occurrence analysis also fails to account for directional interactions and asymmetry like amensalism and commensalism. Some limitations of co-occurrence-based strategies for cross-sectional datasets can be improved by conducting time-lagged correlation of longitudinal observations. In time-lagged correlation, a positive relationship is denoted when the relative abundance of one species at time t correlates with an increase in relative abundance a second species at time t + 1 , and a negative (competitive) relationship is inferred from a correlation with the decrease in the relative abundance of the second species at time t + 1 (14). Time-lagged correlations account for directionality and asymmetry in putative interactions which allow for the modeling of amensal (0/-), commensal (0/+), and exploitative (+/-) relationships between bacteria. Although time-lagged correlation methods are promising, these methods have not been broadly applied to understand microbial relationships in different contexts, including in the context of microbiome disturbance, and have not been coupled with functional information available from databases of annotated genomes and microbial metabolic networks to make molecularly-informed hypotheses about the mechanistic basis of these relationships. To address this gap, we further developed time-lagged correlation methods and applied them to a dense longitudinal study of four Westerners traveling abroad in Bangladesh who experienced diarrhea. This observational study’s high temporal resolution and multi-week time span enabled this analytic strategy. We combine information about these associations with their imputed genomes, metabolic network analysis, and extreme gradient-boosted random forest classifiers to predict the basis of potential cooperative and competitive relationships among gut microbes.

## RESULTS

### Community-level trends and dynamics

Our longitudinal dataset consists of four adult Westerners, three males and one female, who were traveling abroad in Southeast Asia for 2-4 weeks. All 4 individuals lived in Colorado, USA and self-sampled every stool that they produced for 2 weeks prior to and then during and after travel to Dhaka, Bangladesh. Subject F01 was in Dhaka for 1 week and then in Cambodia for an additional week before returning to Colorado, where she continued to collect samples for an additional 2 weeks. M01 and M02 were in Dhaka for 2 weeks before returning to Colorado; M03 was in Dhaka for 1 week before traveling to Israel for a day, Canada for 3 days, and then returning to Colorado. Each individual collected fecal samples at the time of each bowel movement by swabbing used toilet paper, and the samples were subjected to 16S rDNA targeted sequencing.

While in Dhaka, Bangladesh all 4 individuals experienced diarrhea and/or vomiting from a likely food-borne pathogen. The timing of the illness and shared food sources suggested that F01, M02, and M03 all had a shared exposure. The onset of illness for M01 was delayed so may have been from a different exposure. The symptoms and severity of illness varied across the 4 individuals, for instance with F01 experiencing vomiting and relatively mild diarrhea and M01 having severe diarrhea, having collected 13 unique stool samples on the day of illness onset. Examination of the 16S rRNA amplicons did not reveal any compositional blooms of likely bacterial culprits, such as Enterobacteraceae (which would include *Escherichia coli* and *Salmonella* spp.), or *Campylobacter*. During this diarrheal episode, designated by the vertical red bars in Figure 1, subjects M01, M02 and M03 experienced an abrupt decline in the richness of bacteria in the gut as assessed with the phylogenetic diversity (PD) (15) alpha diversity measure (Figure 1A). Despite this drop in PD there were no obvious signs of canonical low- diversity dysbiosis defined as low alpha diversity coupled with increased colonization of facultative anaerobic bacterial families such as *Enterobacteriaceae* or *Lactobacillaceae* (16). Subject F01 had no drop in alpha diversity. Despite a suspected shared exposure in 3 of the 4 individuals, there was no compositional convergence among individuals during illness or recovery, consistent with interpersonal variation being strong in human studies (17). However, the drop in PD did correspond with an increase in turnover within individuals, as measured by Weighted UniFrac distances to samples at the subsequent timepoints (Figure 1D). For the three males, alpha diversity and diarrheal severity, as estimated by the number of stool samples collected per day (Figure 1B), were predictive of community stability with decreased phylogenetic diversity being indicative of increased turnover (*p = 1 x 10^-31^, R^2^ = 0.13*) (Figure S1). All four subjects had *Bacteroides*-dominant microbiomes typical of industrial societies at the outset of their travels, and this enteric disturbance resulted in an enterotype switch for subject M03 wherein *Bacteroides* genera were largely displaced by *Prevotella* (Figure 1C) which is an enterotype canonically associated with agrarian societies or high fiber diets in Western individuals (18). As subject M03 underwent dramatic compositional shifts in his core microbiome which were incompatible with our filtering strategies, we excluded him from downstream analysis involving the identification of putative ecological interactions which we here-in refer to as associations.

**Figure 1:**
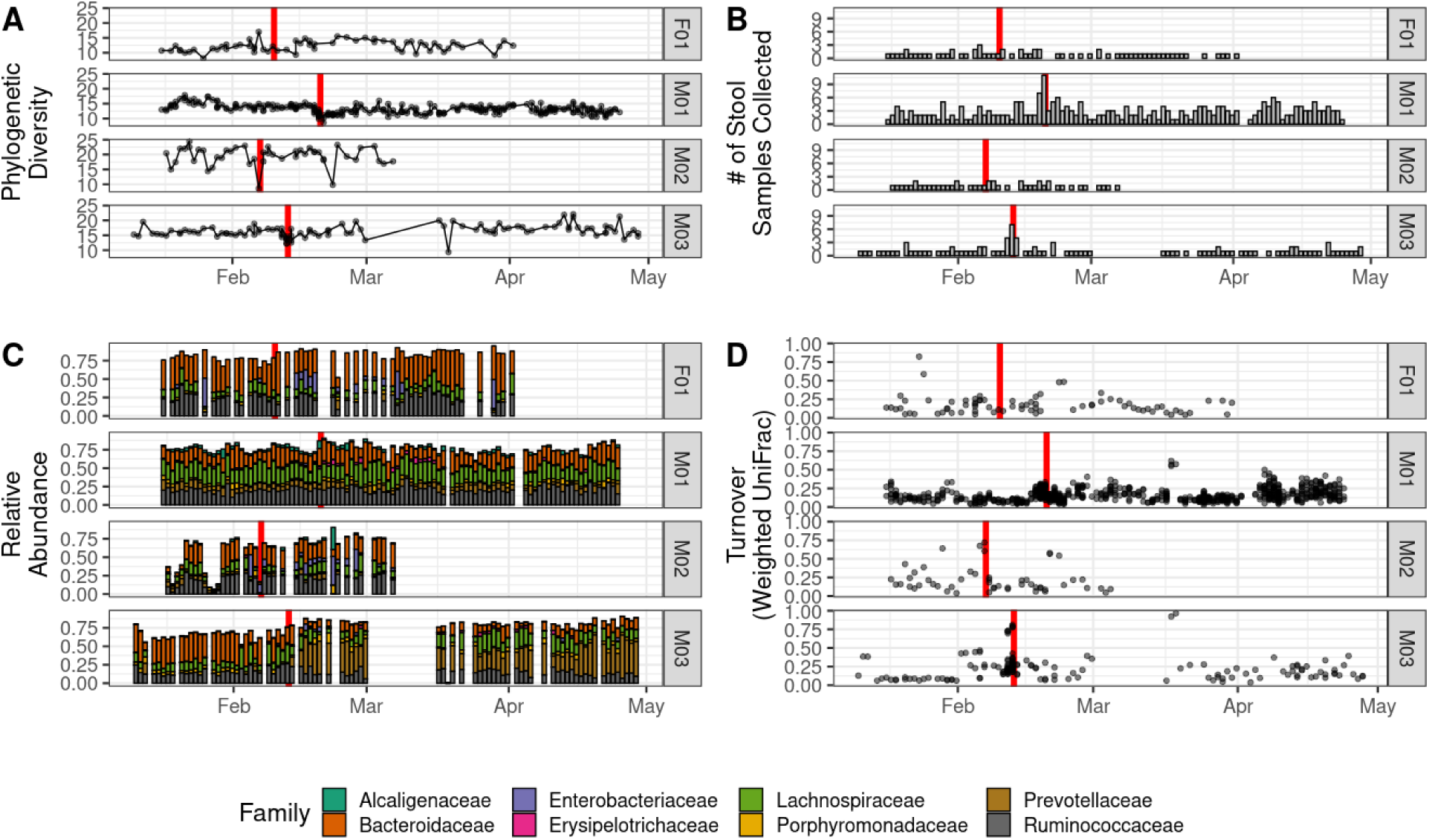
Fecal microbiome composition over time in 4 travelers who experienced diarrhea. Data is from one female (F01) and three males (M01, M02, M03) who experienced diarrhea while traveling in Bangladesh. The red line indicates the timing of the diarrheal episode. A) Phylogenetic Diversity (PD) over time in all 4 individuals. B) The number of stool samples collected per day, showing a peak in M01 and M03 with diarrhea. C) Taxonomic composition (family level) over time. D) Turnover expressed as the Weighted UniFrac distance between each timepoint and the subsequent time point.

### Identification of putative ecological interactions (associations)

To identify associations between microbial relative abundances, we adapted the time-lagged correlation strategy described by Trosvik *et al* (14), and applied it to 97% identity (ID) Operational Taxonomic Units (OTUs). We chose to bin highly related Amplicon Sequence Variants (ASVs) selected with DADA2(19) into 97% ID OTUs, because this threshold has typically been used in microbiome data analyses to bin sequences derived from closely related organisms and to approximate the species level (20). Binning to 97% ID OTUs reduced the sparsity of the data matrix and increased the number of features that were observed commonly across a given individual without impacting resolution since 16S rRNA cannot typically define organisms or impute functional information at the strain level. Within each individual, and for all pairs of 97% ID OTUs (i, j) that were observed in at least 90% of the samples for that individual, we determined OTU pairs for which the centralized log ratio (CLR)-transformed value of OTU j at time t correlated with the change in the CLR-transformed value of OTU i between time t and time t + 1 (see methods). Missing data was imputed, and samples merged such that there was exactly one sample value per day (detailed in methods). A CLR-transformation was applied since compositionality has been shown to impact time-lagged analyses (21) and the CLR-transformation has been shown to be effective at correcting for compositionality-driven artifacts in simulations of non-time lagged correlations detections between OTUs (12). A positive Spearman coefficient β:

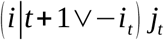

would indicate that higher levels of OTU *j* (relative to the mean) meant greater increases in OTU *i* in the next time point, indicating that OTU *j* may facilitate success of OTU *i*. A negative β would indicate an antagonistic or competitive effect of OTU *j* on *i.* To determine whether β was more positive or negative than expected by chance, we permuted the order of the timeseries by shuffling the time (columns) in the data matrix. This created null distributions where the time value was no longer meaningful while maintaining the relative abundances distributions across OTUs at each timepoint. We assessed statistical significance as βij values that had extreme absolute magnitudes as compared to their corresponding null distribution of βij values. Surprisingly, we observed a positive correlation between permuted and empirical Spearman coefficients for all βij (Figure S2). One indicator of the potential driver of this trend is that null Spearman coefficients for i = j interactions (i.e. comparing an OTU to itself) were all among the most negative. This is demonstrated in Figure S3, where we show this phenomenon for a single OTU and across all OTUs in all individuals. A negative time-lagged correlation for randomized i = j interactions suggests that this is a statistical artifact, where OTUs that are highly positively correlated with each other at the same time points, are negatively correlated in time lagged analysis. We believe that this is driven by “regression towards the mean”, where if one sample of a random variable is extreme, the next sampling of the same random variable is likely to be closer to its mean (22). These trends highlight the importance of using our permutation strategy for assessing statistical significance rather than a traditional p-value. Due to the extreme null attributes of these *i = j* outliers, we omitted them from consideration from our analysis. This methodology identified 78 positive and 140 negative associations out of ∼8,500 considered *i←j* pairs (Figure 2C). Seventy-five of the 78 positive associations were found in Male 01 who also had the longest and most densely sampled time series; the remaining three positive associations were found in Female 01. A complete list of associations with their taxonomic assignments and direction of change is provided in Supplemental Table 1.

**Figure 2:**
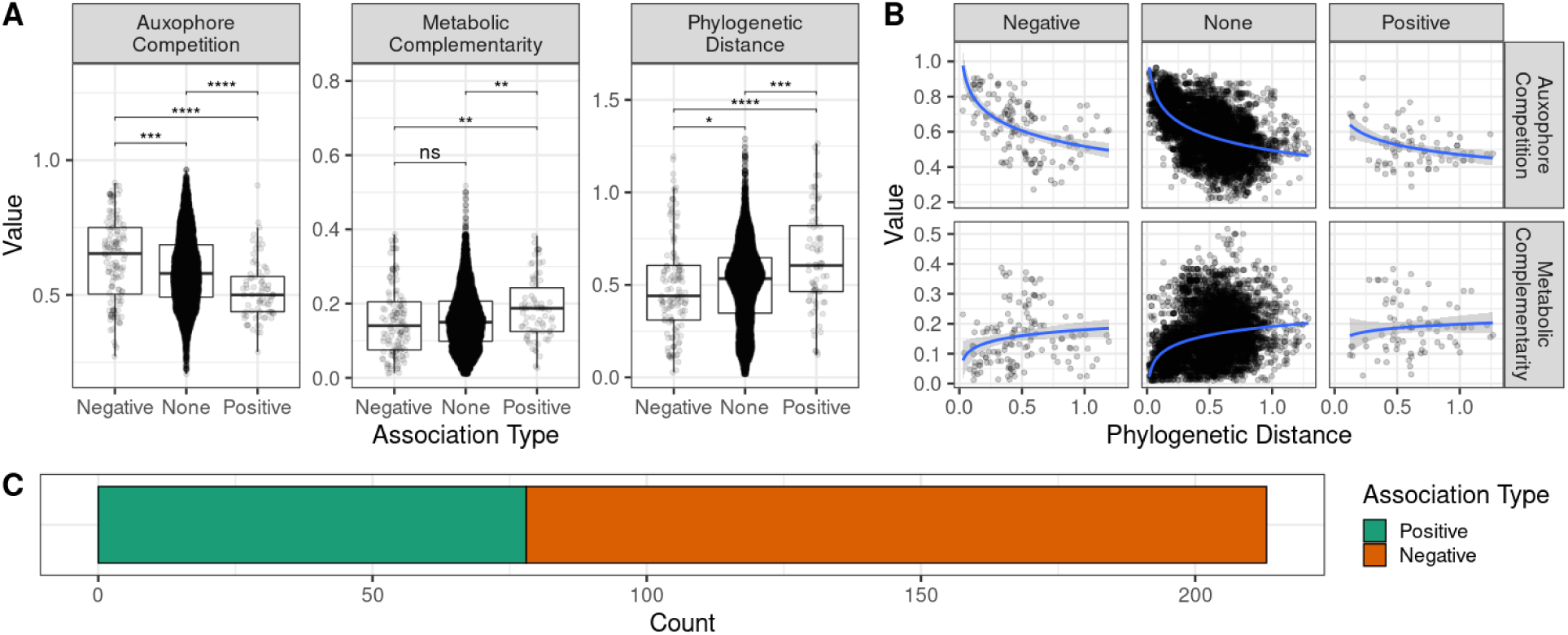
Phylogenetic and functional patterns across association types. A) Estimated rates of auxophore competition and metabolic complementarity by association type as calculated by NetCooperate. **\*** p < 0.05, **\*\*** p < 0.01, **\*\*\*** p < 0.001, **\*\*\*\*** p < 0.0001. B) Estimated rates of auxophore competition and metabolic complementarity as a function of phylogenetic distance and association type. The curves for the positive associations are different from those of the negative and null associations (p < 0.05); there is no difference between the negative and null association curves. C) Relative fraction of positive (78) to negative (140) associations.

We found that the positive associations were between bacteria that were significantly more phylogenetically distant than the negative and null *i←j* pairs and that the negative associations were between more closely related bacteria (Figure 2A). As bacterial metabolic niche space is roughly correlated with phylogeny (23), we hypothesized that these patterns in relatedness across association types were a reflection of metabolic competition and cooperation. To test this hypothesis, we used PICRUSt (24) to impute bacterial genomes and then mapped the available enzymatic reactions to KEGG (25) in order to generate a putative metabolic network for each 16S rDNA amplicon. We then used these metabolic networks as inputs for NetCooperate (26), which returns information about the relative metabolic potential between two bacteria’s respective metabolic networks. NetCooperate estimates metabolic complementarity as the fraction of auxophores (27) (essential substrates which cannot be prototrophically synthesized) required by OTU *i* which can be synthesized by OTU *j*. We expanded NetCooperate to measure metabolic niche overlap, which is defined as the fraction of OTU *i*’s auxophore pool that is also required by OTU *j*. We identified higher potential for metabolic cooperation amongst the positive associations compared to both the negative and null associations (Kruskal Wallis test with Dunn’s post-hoc test; p < 0.01), but there were no differences between the negative and null groups (Figure 2A). We found substantial differences in auxophore competition between all association types with the negative associations having the highest metabolic niche overlap followed by the null, and positive associations (Figure 2A). Together, these observations indicate that negative associations are enriched for microbe pairs that are more phylogenetically related, have lower potential for cooperation and a high degree of niche overlap, and the inverse is true for positive associations. To test if these metabolic relationships were merely a function of phylogeny and niche space co-correlation, we modeled these measures as a function of phylogenetic distance with a statistical interaction with the corresponding association type (negative, null, and positive). We observed that when adjusting for phylogenetic distance, positive associations still had higher metabolic cooperative potential and lower niche overlap compared to the negative and null associations, however, there were no differences between the negative and null association’s curves (Figure 2B).

To find specific metabolic signatures that could provide mechanistic clues for the basis of these potential ecological interactions, we searched through the compounds that NetCooperate deemed to be metabolically complementary or competitive for each pair of associated OTUs. We identified substrates that were required by both organisms and which were specifically enriched in negative associations but not in positive ones using a Z-score threshold of two. Using this approach, we found three different compounds: 1) glyoxylate, a two-carbon carboxylic acid, 2) myo-inositol, a carbocyclic acid, and 3) nicotinate, vitamin B3 (Figure 3a). We used a similar approach to find metabolically complementary compounds that were enriched in positive associations yielding seventeen different compounds. Broadly speaking, these 17 metabolites can be categorized as fatty-acid carriers, nucleoside/nucleotide derivatives, and carbohydrates (Figure 3b).

**Figure 3:**
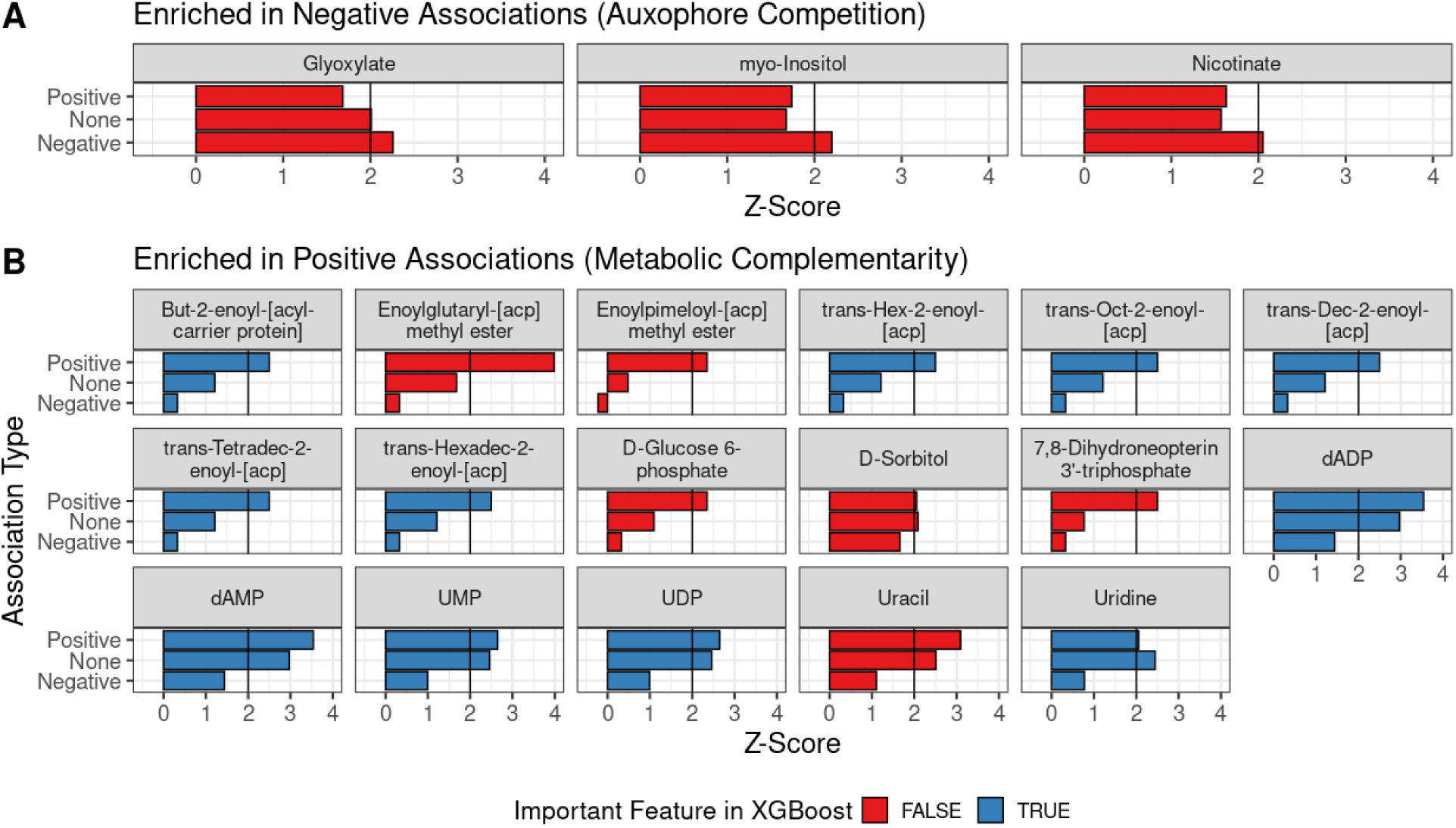
Compounds identified using NetCooperate that were significantly enriched in Negative or Positive associations. A) Metabolically competitive auxophores that were enriched in negative but not positive interactions compared to pairs with no detected relationship using a Z-score threshold of 2. B) Metabolically complementary compounds enriched in positive but not negative interactions using a Z-score threshold of 2. Plots are colored blue if the compound is a substrate of KEGG Orthologies (KOs) determined to be important by a trained extreme gradient boosted random forest classifier (XGBoost) - see Figure 4.

Though we were able to detect significant differences between the positive and negative association types in phylogenetic distance, metabolic complementarity, and auxophore competition, logistic classifiers that used these univariate summary measures were incapable of distinguishing unlabeled associations (Figure 4A). We also tested trait similarity, by forming vectors of both enzymatic and non-enzymatic orthologs using assignments to the KEGG Orthology (KO) database that were made from each OTUs genome predictions by PICRUSt. We then calculated the Jaccard distance of the two trait vectors (a list of given KOs) for each OTU pair. We found that this raw trait similarity measure was not predictive of the association type when using a simple logistic regression *β_ij_ J* (*^i^*^⃗^, ^⃗^*j* ) (Figure 4A). We hypothesized that since the NetCooperate analysis identified metabolites that differed between positive and negative association types, a high dimensional feature set that explicitly included trait data from both OTU *i* and *j* instead of a summary difference/similarity metric would provide more discriminatory power. To this end, we trained an extreme gradient-boosted random forest classifier (28) (XGBoost) to predict the association type using the concatenated trait vectors of OTUs *i* and *j*.

**Figure 4:**
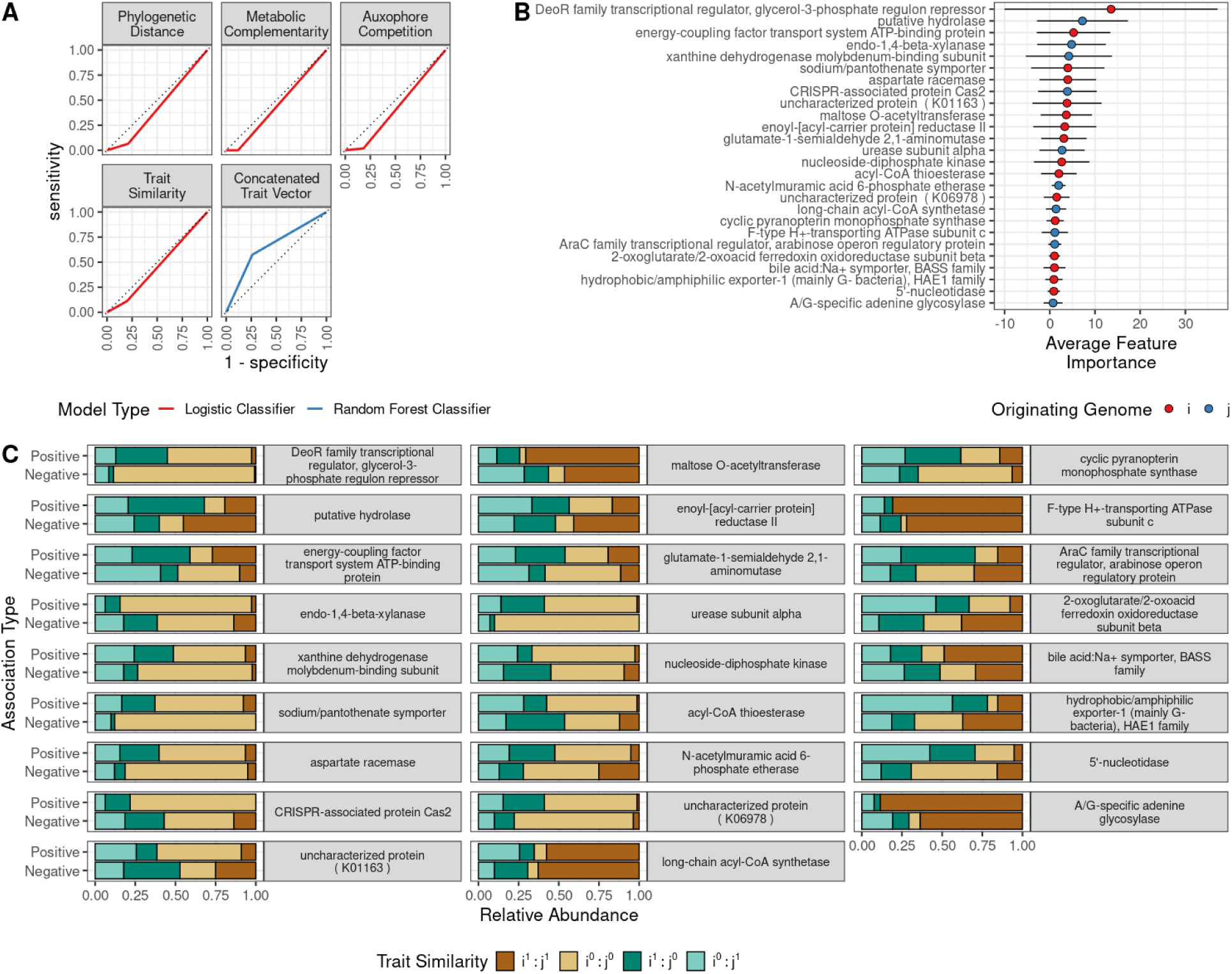
Extreme Gradient-Boosted Random Forests (XGBoost) trained on concatenated KO presence/absence trait vectors can discriminate between positive and negative associations from time-lagged correlation. A) Receiver Operator Characteristic (ROC) curves demonstrating that Random Forest classifiers trained on high-dimensional trait vectors outperform logistic classifiers that use univariate summary measures. B) Informative traits as determined by permutation testing using BorutaShap. Features are colored by which OTU’s genome (*i* or *j*) the trait originated from. C) Trait concordance across informative traits found by XGBoost + BorutaShap. (*i^1^ : j^1^* and *i^0^ : j^0^*) - trait is present or absent in both organisms. (*i^1^ : j^0^*) - Trait is present in *i* and absent in *j*. (*i^0^ : j^1^*) - Trait is absent in *i* and present in *j*.

In this case, the trait vector is a bit vector of 1’s and 0’s indicating presence or absence of PICRUSt-inferred KOs. For instance, for each inferred KO and each *i←j* pair of OTUs, we encoded (0,0) if absent in both, (1, 1) if present in both, and (0,1) or (1,0) if present in one and not the other. We found that an XGBoost classifier achieved ∼72% classification accuracy (Wilcoxon test; p < 0.001) as determined by 5-fold cross-validation, and the receiver operator characteristic (ROC) curves substantially outperformed the univariate measures (Figure 4A). We trained the XGBoost classifier using the estimated KO copy number as predicted by PICRUSt as well as simple presence/absence information for each KO, and we detected no differences in performance between the models. As a result, we used the KO presence/absence trait vectors as they were simpler to interpret. We afterwards used BorutaShap (29), which uses permutation to determine whether features have a higher importance score than chance expectation, at a threshold of 0.9 on the trained XGBoost model, to extract important KOs for classifying association type, and identified 26 PICRUSt inferred KOs with discriminative power (Figure 4B). Sixteen of these informative KOs were from OTU *i’s* genome while the other ten originated from OTU *j* (Figure 4B). Notably, 65% (11) of compounds that we identified as enriched in positive associations in the metabolic complementarity measures (Figure 3B) are also substrates of three enzymatic KOs that BorutaShap deemed to be important in the XGBoost classifier. The various enoyl-acyl carriers (Figure 3B) are potential substrates of enoyl- [acyl-carrier-protein] reductase II (K02371) while the nucleoside/nucleotide derivatives are substrates of the 5’-nucleotidase (K01081) or nucleoside-diphosphate kinase (K00940). There are four different states for every trait in a given *β_ij_* pair: the trait is present or absent in both organisms (*i^1^ : j^1^* and *i^0^ : j^0^*), or the trait is present in *i* but not *j* (*i^1^ : j^0^*) or vice versa (*i^0^ : j^1^*). We compared the relative frequencies of these relationships and found that trait discordance (*i^1^ : j^0^*and *i^0^ : j^1^*) was more common in positive associations than in negative ones (p < 0.05). We illustrate the general patterns of informative trait discordance, given in shades of green, and concordance, given in shades of brown, in Figure 4C.

We next visualized the state space of the 26 informative KOs identified using XGBoost + BorutaShap by embedding each OTU involved in an association into a principal component analysis (PCA) space based on the Jaccard distances between their presence/absence data for the 26 KOs (Figure 5A). The first two principal components accounted for ∼62% of the variance; PC1 linearly separates two primary clusters: 1) a dispersed grouping of *Bacteroidales* (*Bacteroides*, *Parabacteroides, Prevotella,* and *Paraprevotella*) with *Desulfovibrionales* (*Bilophila*) and *Burkholderiales* (*Sutterella*) on the left and 2) various Clostridiales species, *Coriobacteriales*, and an assigned member of *Bacteroidaceae* on the right. We observed that the positive associations occurred predominantly between these two clusters, and negative associations were more common within clusters. We also illustrate the putative presence/absence of select traits (5’ nucleotidase, enoyl-acp-reductase II and a sodium/pantothenate transporter) that independently appeared in both the metabolic network analysis and the XGBoost + BorutaShap feature selection.

**Figure 5:**
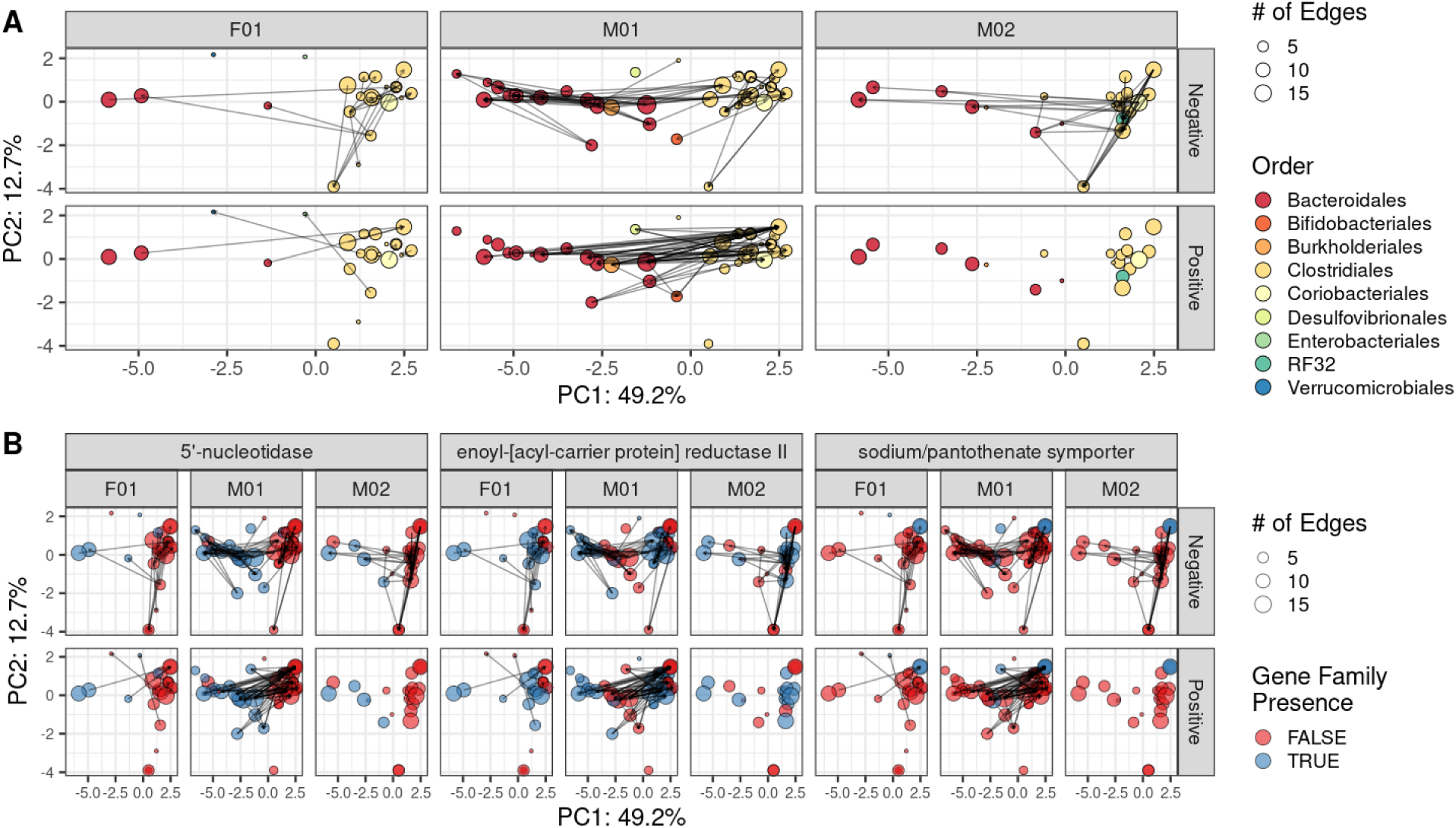
OTU Principal Component Analysis (PCoA) space using KOs identified by XGBoost + BorutaShap. A) OTUs, given as nodes colored by taxonomic order, that are closer in PCoA space are more functionally similar. Negative (top row) and positive associations (bottom row) are depicted as arrows between nodes (*j* → *i*). Columns designate the observed associations for the three individuals. B) Gene family presence absence data for 5’-nucleotidase (K01081), enoyl acyl-carrier protein reductase II (K02371), and sodium/pantothenate symporter (K14392). Red indicates an absence of the gene family for a given OTU and blue is presence.

## DISCUSSION

In this study, we applied time-lagged correlation to dense time series data collected from travelers who had diarrhea to predict taxa pairs that had positive or negative associations indicating potential cooperative/facilitative or competitive relationships, respectively. We then used PICRUSt predictions of genomic content, metabolic network modeling and random forest to identify inferred metabolites and genes that predicted these relationships, generating novel hypotheses regarding the underlying driving factors of microbe-microbe interactions in the human gut. This work has thus produced a rich collection of hypotheses regarding interactions between microbes that could be tested in the lab. Understanding specific facilitative and antagonistic relationships that influence the success of microbes in the gut will facilitate efforts to modify microbiome composition to promote health.

In this time series, the four subjects were suspected to have had a foodborne exposure to an enteric pathogen. Our search through the 16S rRNA amplicons did not identify a plausible bacterial source, suggesting an alternate etiology, such as a viral pathogen. We observed a sharp drop in phylogenetic diversity at the time of enteric disturbance in Males 01, 02, and 03 (Figure 1A) but did not detect convergence to a common microbiome composition. This illness was accompanied by decreased community stability, and in the case of subject M03, a drastic enterotype conversion to a *Prevotella*-rich microbiome composition. Changes in diet and water source can have a profound impact on microbiome structure and function, and immigrants from Southeast Asia to the United States experience distinctive shifts from a *Prevotella-*dominant to *Bacteroides*-dominant composition (18). We hypothesize that Male 03’s diarrheal event depleted community richness and opened new gastrointestinal niches, and the change in diet and environmental exposures due to travel in Southeast Asia led to a concordant enterotype conversion. We also found that the community dynamics (Figure S1) were consistent with the theory that high microbiome diversity begets community stability (lower turnover), however, the strength of this relationship was specific to the individual and only accounted for 13% of the variance. Applying our time lagged correlation analysis to microbiomes that underwent disturbance and remodeling should in principle have increased the number and type of interactions observed since repeated measurements of a more stable microbiome would not be expected to contribute as many highly informative data points (30). Since microbiomes do typically vary somewhat over time due to factors such as normal dietary variation or exposures, further studies with time series data collected with and without disturbance would be needed to determine the importance of disturbance in identifying relationships.

Co-occurrence has often been used to infer ecological interactions, and its use and interpretation have been a matter of debate for over a century (13). Analysis of co-occurrence and microbial recruitment patterns in human microbiomes has indicated that co-occurring bacteria tend to be more closely related than chance expectation, and that this phenomenon is likely a signature of environmental filtering rather than direct interactions (10, 31). The time-lagged-correlation strategy used here was applied previously to time series data collected from 2 individuals (14). One notable difference in our study was that we used a permutation-based method to determine statistical significance that corrected for underlying bias in each βij microbe pair. Both we and Trosvik *et al* found that empirical βij where *i = j,* had a large negative Spearman coefficient, and Trosvik et al suggested that this was due to abundances of taxa over time being influenced by the carrying capacity of an environment for that taxon. However, our finding that the null βij distribution was more negative than empirical βij where *i = j*, suggests an underlying statistical artifact rather than a biological explanation. We instead hypothesize that this behavior an example of a “regression to the mean” which is a phenomenon where a randomly sampled observation with an extreme value is more likely to be followed by random sample with a value closer to the population mean. Our finding that | *null* βij | > | empirical βij | indicates that there is actually less autocorrelation of taxa with themselves than you would expect by chance, perhaps suggesting factors against species turnover within communities. The overall positive correlation of empirical βij with randomized βij values would derive from pairs of *i* and *j* showing regression to the mean effects at a strength related to the degree of correlation. Future studies employing time-lagged correlations should thus use our permutational strategy for assessing statistical significance to avoid false positives. Note that other published microbiome association methods such as Local Similarity Analysis (LSA) (32) have used permutation strategies that have been suggested to sometimes produce false positives (33). However, these prior analyses have permuted *i* and *j* independently of each other rather than preserving the OTU relative abundance distributions at each time as we do here (i.e. shuffling the order of each row in a feature by time matrix rather than order of the columns). However, other methods developed to compute a null distribution for correlation that do not destroy autocorrelation structure in the data have been developed and would be interesting to also apply, including the IAAFT (iterative amplitude-adjusted Fourier transform) method and the Twin method (34) and other parametric approaches (35, 36).

We used a correlation strategy as opposed to a regression or dynamic systems model, which can have certain advantages over correlation, including the ability to control for confounders, account for perturbation frameworks, use data from multiple individuals together while controlling for dependency (rather than stratifying by individual), and forecast future system behaviors(30). Regression based strategies may also be influenced by the regression to the mean artifact described above and this should be investigated further. Although sophisticated generalized Lotka-Volterra (gLV) based models have been developed and shown to be effective, such as MDSINE(30),TPG-CODA(38), and compositional Lotka-Volterra (cLV) (21), these have only been used to model a small number of taxa and have analytical challenges to scale to whole human microbiomes(30). The linear mixed model based approach of MTV-LMM has been run at scale but was developed to identify taxa whose temporal dynamics depend on whole community composition rather than the pairwise interactions between individual taxa (37). Other methods such as the sparse vector autoregression (sVAR) model (33), have been applied to large datasets and used to differentiate autoregressive OTUs whose abundance dynamics depend on community composition at previous timepoints, as well as non-autoregressive OTUS whose abundance was more related to changes in external factors such as diet. With the sVAR approach, time-lagged interactions between OTUs within the model could be estimated using autoregressive model coefficients equivalent to partial Granger coefficients (39), which could be a promising alternate approach to apply here. Interestingly, prior work that applied regression models to time series data have found that a considerable portion of the human gut microbiome have time-dependent relative abundance patterns that can be predicted by microbiome composition (30, 37) and have recovered experimentally-supported relationships between microbes in smaller synthetic communities evaluated over time in gnotobiotic mice, such as an inhibitory effect of *Clostridium scindens* on *Clostridioides difficile* (*30, 40*).

We found a higher prevalence of negative associations between phylogenetically similar OTUs, suggesting an importance of competitive interactions in shaping gut microbiome composition over time, but we also observed a high number of positive associations indicating potential cooperative relationships. The ratio between cooperative and competitive relationships in ecological networks and the resulting impact on microbiome function, stability, and resilience is currently uncertain; mathematical frameworks have produced seemingly conflicting conclusions that both higher and lower ratios of competition:facilitation can reduce community resistance to ecological perturbation (4, 41). Moreover, interaction strength and the interaction network topology are also conjectured to influence community-level attributes (41, 42). In the mammalian gut, both competitive and cooperative relationships have been demonstrated to augment community resilience and stability, and the successful explanatory models will almost certainly need to incorporate contextual details like network topology, spatial architecture, and temporal variability (43). The significance for competition was further supported in our analyses by the lower levels of shared auxophores based on the predicted metabolic networks of positively associated pairs compared to the null and negative pairs. This result is consistent with experimental and mathematical models of the mammalian gut microbiome that have depicted an ecosystem with prevalent competition for limited resources (7, 42, 44–46).

Our analyses also resulted in the detection of many positive associations that had higher predicted metabolic complementarity, supporting that these cooperative relationships also play an important role in community dynamics over time. This is consistent with prior studies that have linked high alpha diversity and CR with complex cooperative cross-feeding in the microbiome (5, 47, 48). Metabolic cooperation has been previously documented for substantial carbon sources in the gut, especially at the epithelial mucosa where microbes collectively hydrolyze large, branched, host-derived glycoproteins into extracellular “public goods”(8, 9). Sharma *et al* also observed a high prevalence of division of labor and cooperative micronutrient salvage within the gut (8). They noted that auxotrophy for major B-vitamins was common throughout the bacterial community and showed that there are essential, committed B-vitamin salvage pathways between commensal bacteria. Moreover, they demonstrated that loss of the cooperative relationships between various B-vitamin prototrophs and auxotrophs destabilized the overall microbiome composition, indicating that there are likely numerous other interaction-mediated influences on community structure and function.

Our NetCooperate metabolic network analysis and XGBoost + BorutaShap model independently found three distinct classes of molecules whose predicted levels varied between positive and negative associations: 1) B vitamins pantothenate and nicotinate, 2) enoyl acyl-carrier protein (acp) reductase II (FabK) and 3) nucleotide/nucleoside derivatives. NetCooperate highlighted mutual nicotinate auxotrophy in negative associations while XGBoost + BorutaShap identified dissimilarity (*i^1^:j^0^* and *i^0^:j^1^)* in 5’-nucleotidase (K01081), which is involved in the metabolism of nicotinate-nucleosides, as being enriched in positive associations. These findings suggest competition and metabolic niche partitioning regarding nicotinate (B3) synthesis. Magnúsdóttir *et al* conducted genomic queries for various B vitamin auxotrophies across gut commensals, and they observed that environmental nicotinate salvage was unique to members of *Actinobacteria*, *Firmicutes*, and a single *Proteobacteria* (49). We detected numerous negative associations among members of these phyla (Figure 5B), and we hypothesize that these relationships may be influenced by competition over extracellular B3 pools. Our feature selection approach also identified the sodium/pantothenate (B5) symporter as an informative feature: Specifically, when OTU *i,* the microbe being influenced by *j*, was predicted to have the pantothenate symporter, *j* was not (*i^1^:j^0^)*. Here, we hypothesize that *j* is likely a B5 prototroph as it does not have a transporter, and *j* donates B5 to *i,* which takes up the “public good” via the sodium/pantothenate symporter. *In vitro* B vitamin cross-feeding assays using *Escherichia coli* demonstrate that B5 prototrophs are indeed able to donate vitamins to co-cultured auxotrophs and strongly rescue growth in a B5-deficient medium; however, this group also noted high variance between differing donor strains which suggests additional mediating factors in cross-feeding efficiency (49). As both B5 and B3 are severely depleted in ulcerative colitis, ecological relationships governed by B vitamin dynamics may represent a therapeutic target for future clinical interventions (50). Enoyl acp reductase II (FabK) catalyzes the terminal step of fatty acid elongation and is widespread in Clostridiales members (51). We were unable to find any literature regarding enoyl acp reductase involvement in bacterial interactions; however, *in vitro* experiments on Δ*fabK*Δ*fabI Enterococcus faecalis* strains revealed that fatty-acid deficient bacteria can be cultured in the presence of exogenous fatty acids (52).

Although our study produced very intriguing and interpretable results, we acknowledge weaknesses. One is that functional genes and metabolites were made based on 2 levels of inference 1) PICRUSt was used to infer genes present from 16S rRNA and 2) NetCooperate was used to infer metabolite auxophores based on these genes. PICRUSt does not predict genes that are subject to frequent horizontal gene transfer, and we may have missed interactions that are based on genes/metabolites that are subject to high strain level variation or that were among OTUs not well represented in genome databases. Further study with dense shotgun metagenomic sequence data would be beneficial, but given the large number of samples needed for this type of analysis, are often prohibitively expensive. Follow-up studies to verify gene presence/absence calls and targeted metabolomics to survey levels of metabolites predicted to be important could help further validate and refine predictions. Correlation-based inferences of interactions have been shown to perform poorly in certain contexts such as in predicting simulated interactions between viruses and bacteria (54), but as applied here, these inferences were able to identify positively and negatively interacting pairs with independently determined metabolic signatures consistent with cooperative and competitive interactions, respectively. In summation, we find that our methodology corrects for spurious associations by using permutation-based testing. The downstream metabolic network analysis coupled with interpretable machine-learning approaches identified distinct functional signals across positive and negative relationships and fostered the generation of targeted hypotheses about the mechanistic basis of the relationship. Our findings support that both competitive and cooperative relationships shape gut microbiome compositional dynamics over time.

## MATERIALS AND METHODS

### Sample Collection

Fecal samples were collected at the time of each bowel movement by swabbing used toilet paper (55). Swabs were stored at room temperature during travel and at - 80^ο^ C upon return and prior to 16S rRNA targeted sequencing. The human volunteers gave consent under the University of Colorado, Boulder, IRB protocol 0409.13.

### Sample processing

DNA was extracted using the MP Biomedicals Powersoil kit following the Earth Microbiome Project (EMP) protocol (http://www.earthmicrobiome.org) (56). Barcoded primers targeting the V4 region of 16S rRNA were used to PCR amplify the extracted bacterial DNA also using the EMP standard protocols. Quantification of PCR products was completed using PicoGreen (Invitrogen, Carlsbad, CA). The UltraClean PCR Clean-Up Kit (MoBio, Carlsbad, CA) was used to clean and pool equal amounts of DNA from each sample. Sequencing was conducted using a MiSeq personal sequencer (Illumina, San Diego, CA).

### Sequence Processing

Samples were demultiplexed using QIIME 1 (57), trimmed to a sequence length of 94 base-pairs, and denoised via Dada2 with the recommended default settings for maximum number of N’s in the amplicon sequence (maxN = 0), truncation quality (truncQ = 2), phiX removal (rm.phix = TRUE), and the maximum number of expected errors in a read (maxEE = c(2,2)). The resulting amplicon sequence variants (ASVs) were then binned into 97% OTUs using UCLUST’s (58) closed-reference schema. Sequences were aligned using pyNAST (59) and the <u>core_set_aligned.fasta.imputed</u> reference alignment downloaded from greengenes.lbl.gov. Fasttree (60) was used to create a phylogenetic tree using the <u>lanemask_in_1s_and_0s</u> file from greengenes.lbl.gov to mask highly variable positions in the alignment. Samples were rarefied to 7500 sequences per sample. Taxonomic assignments were made using greengenes 13_8. Alpha diversity (PD) and beta diversity (Weighted UniFrac) were calculated using QIIME 1.

### Lag-Correlation

We performed CLR transformations on relative abundance data for each of our samples to account for the compositional nature of microbiome data, which has been shown to have the potential to impact time-series analyses, particularly in cases such as this when absolute abundances of microbes may change over time due to a perturbation (21, 30, 61). We then removed OTUs which were not present in at least 90% of the timepoints for a given individual in order t<u>o</u> focus our analysis on the most prevalent taxa and reduce loss of power while correcting for multiple comparisons. Due to the high variability in composition and thus many of the OTUs were present in < 90% of timepoints, M03 was excluded from downstream analyses. Having equal spacing between timepoints is important in time-series data because the degree of change can be influenced by the length of time between samples(61, 62). To produce a dataset with only one sample per day, we first created a representative sample for days with multiple samples by calculating the median CLR abundance for each OTU. To infer abundances for days where no bowel movements occurred, we used scipy’s Piecewise Cubic Hermite Interpolating Polynomial (PCHIP) (63). We chose this method because it maintains abundances above zero, preserves monotonicity, and avoids overshooting in cases of non-smooth data, and is thus well-suited for microbiome data analysis (64), and it has been applied in time-series analyses of microbiome data previously (33). We performed lag-correlation analysis separately for each individual. For each pair of OTUs (i,j), we determined the Spearman rank-correlation coefficient between the CLR-transformed relative abundance of OTU j at time t and the change in the CLR-transformed relative abundance of OTU i between time t and time t + 1. The procedure for generating these coefficients is given as follows:

Let *M be a* 2 x *n* matrix representing a time series of relative abundances of two taxa, *i* and *j*. The rows of *M* correspond to taxa 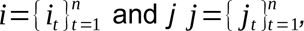, and th*e* columns represent the chronologically ordered time points *t* ∈ {1, 2, 3, … *n*}. The empirically observed Spearman coefficient is calculated as

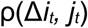

where

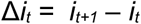

A permutation function, σ, is a bijective function *σ* :{1,2*, … n* }⃗ {1,2 *,… n*} which reorders the column indices of *M* by uniquely mapping each input index to given output index. For example, consider a permutation *σ* (*t* ) for *t* ∈{1 ,2 , 3 , 4 }:

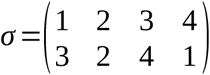

This means:

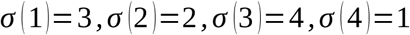

Applying this function to *i* and *j* results in:

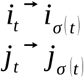

Thus, the permuted Spearman coefficient is then defined as:

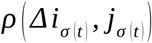

where

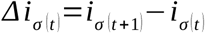

To determine whether calculated Spearman coefficients were more extreme than chance expectation, we compared them to a null distribution which was generated by permuting the time series and recalculating the corresponding Spearman coefficient 2 x 10^6^ times. This method was preferred over traditional p-value calculation for Spearman correlation as applied in Trosvik *et al* (14) because of a strong positive correlation between true Spearman correlations and those calculated based on a null distribution (Figure S2). We used numpy v1.26.0 and scipy v1.11.2. We used the Benjamini-Hochberg FDR correction and filtered for associations with a corrected p-value < 0.1. All statistical calculations downstream of the permutation analysis were made using the R programming language v4.1.2 “Bird Hippy.” Declarative specifications for the analysis environment can be found at https://github.com/casey-martin/bangladesh_time_series.

<u>Genome Imputation (PICRUSt):</u> We used PICRUSt 1’s pipeline as described in their tutorial (https://picrust.github.io/picrust/tutorials/genome_prediction.html#genome-prediction-tutorial) to normalize 16S rRNA copy numbers and impute the corresponding genome contributions for the 97% OTUs and selected Kegg Orthologs as the output trait (24). PICRUSt’s precalculated tables for genome estimation were accessed using the following links: 16S Copy Number Normalization: http://kronos.pharmacology.dal.ca/public_files/picrust/picrust_precalculated_v1.1.4/13_5/16S_13_5_precalculated.tab.gz

GreenGenes 13.5 KEGG Ortholog Table: http://kronos.pharmacology.dal.ca/public_files/picrust/picrust_precalculated_v1.1.4/13_5/ko_13_5_precalculated.tab.gz

### Metabolic Network Analysis (NetCooperate)

The enzymes from the PICRUSt genome predictions were mapped to the KEGG database (25) and approximate metabolic networks were constructed using the archived 2012 KEGG compound database. Each KO was mapped to its associated list of reactions, and then each reaction was split into a directed graph of reactants → products and reactants ← products if the reaction was labeled as bidirectional. In the case where there were multiple reactants or products, all compounds were joined with a directed edge. For example, the formula A + B → C would yield an edge list of (A, C), (B, C). All compounds were then unified into a graph, and subgraphs of size < 10 were removed, leaving only the dominant, contiguous metabolic network for consideration of metabolic overlap and complementarity. NetCooperate provided measures of metabolic complementarity (fraction of auxophores required by OTU *i* which can be synthesized by OTU *j*) were used in downstream analyses. We also estimated metabolic niche overlap (defined as the fraction of OTU *i*’s auxophore pool that is also required by OTU *j*, using auxophore lists output by NetCooperate and custom python and R scripts.

### XGBoost + BorutaShap and PCA visualization

The full genome predictions produced by PICRUSt (both enzymatic and non-enzymatic KOs) were used to train the XGBoost Random Forest Classifiers(28) which consisted of 500 estimators with a max depth of 5. Model performance was not sensitive to these parameters as determined by coarse parameter space search. We tested a range of estimators [500, 1000, 10,000] and max depth [5, 7, 10]. Statistically significant features were identified using BorutaShap’s (29) percentile cutoff of 0.9 over the course of 100 trials. Model performance was estimated using a k-fold cross validation scheme using k = 5. We visualized the state space of 26 informative KOs identified using XGBoost + BorutaShap by embedding each OTU involved in an association into a PCA space. For each OTU, the 26 KOs were encoded as a binary vector, with a one signifying trait presence and a zero signifying trait absence.

## DATA AVAILABILITY

The datasets generated during the current study are available in the ENA repository, https://www.ebi.ac.uk/ena/browser/view/PRJEB69530. Data processing and analysis scripts may be found at: https://github.com/casey-martin/bangladesh_time_series.

## ACKNOWLEDGEMENTS

We would like to thank Luke Ursell for his help in conceptualizing the study and editing, Manuel Lladser for help with mathematical notations and Abigail Armstrong, Laurie Lyon, John Sterrett, and Jack Darcy for the many helpful conversations which ultimately guided the direction of this work.

---

**Figure S1:**
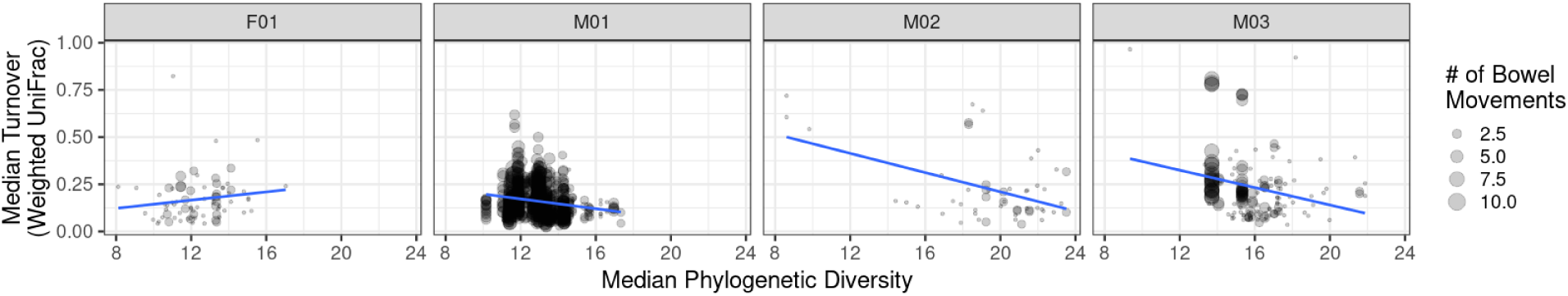
Microbiome stability, measured here as median turnover with the subsequent time point, modeled as a linear mixed effects model of PD + # of Bowel Movements + Individual Effects (p < 0.0001).

**Figure S2:**
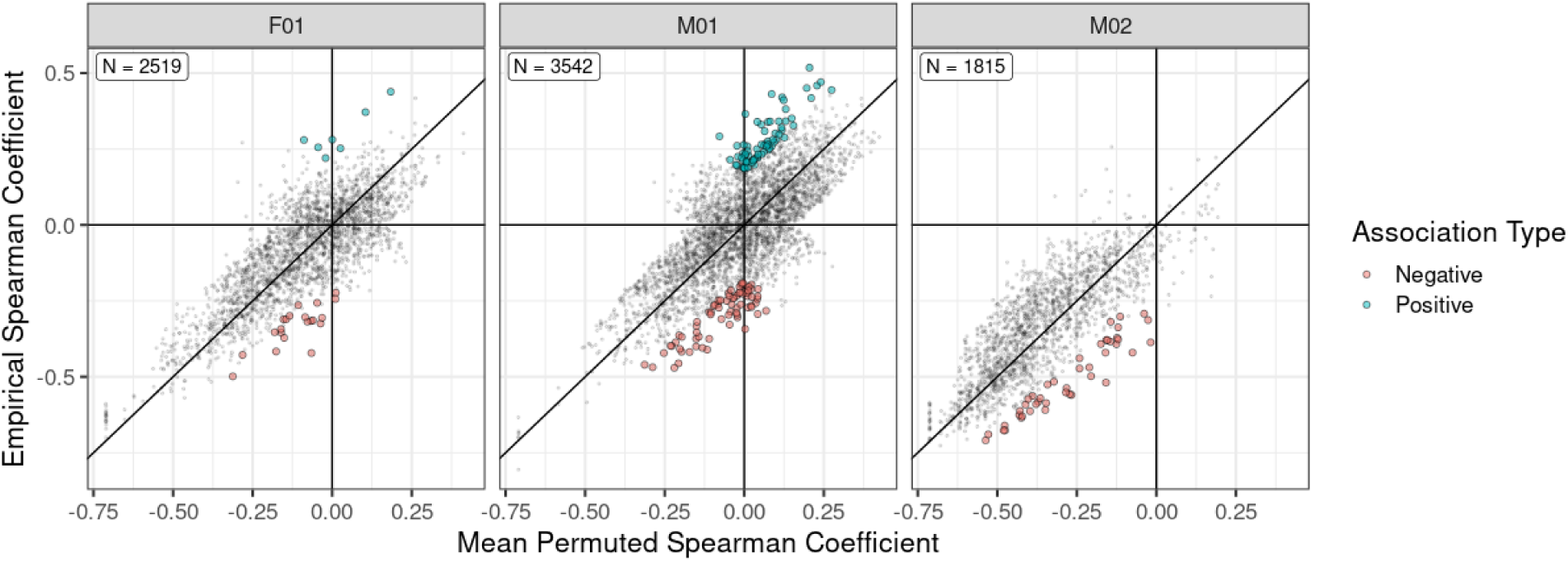
Plot of empirical Spearman correlation R coefficients versus the mean permuted R coefficient, which is calculated based on a null distribution generated by independently permuting the order of the time series and recalculating the Spearman R coefficient 2 x 106 times. Each point represents values for a single OTU i, OTU j pair and all possible pairs are shown. The plot is faceted by whether the OTU pair was observed in F01, M01, or M02. Significant pairs, which are colored by whether the relationship was negative or positive, had an FDR corrected p-value < 0.1. Uncorrected p-values were calculated as the fraction of times that the observed R Coefficient was more extreme than then permuted R coefficients.

**Figure S3:**
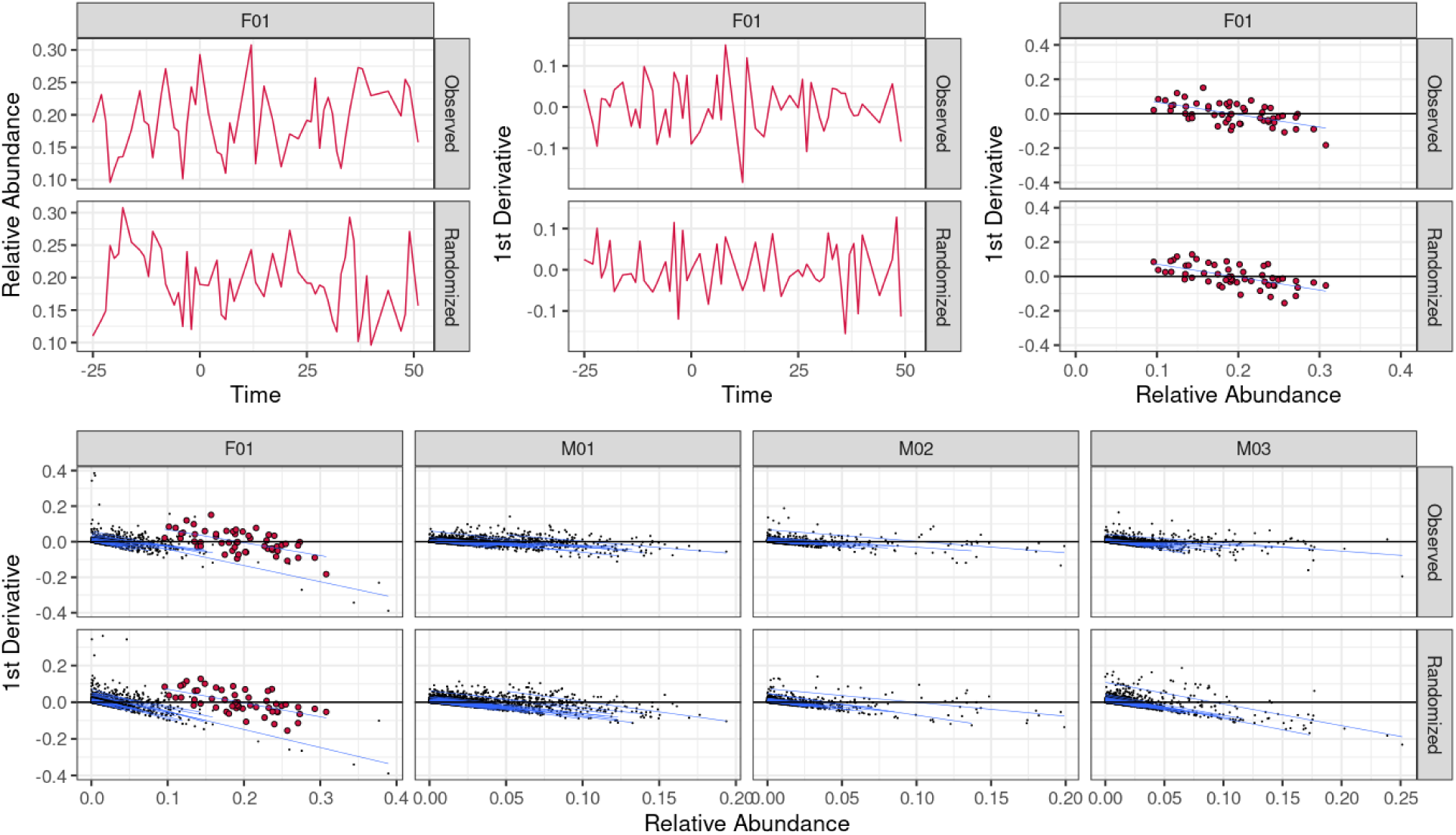
Time-lagged correlations of OTUs with themselves are negative, even in randomized data, suggesting a systemic bias. (A): the observed (top) and randomized (bottom) relative abundance of a single OTU in F01 over time. (B) The 1^st^ derivative ((Δ*it = it+1 – it*) where *i* is the OTU) as calculated from the observed (top) and randomized (bottom) data. (C) Plot of the 1^st^ Derivative of this OTU versus its relative abundance in both the observed (top) and randomized (bottom) data. (D) Plot of the data while comparing all OTUs to themselves in each of the 4 individuals (labeled F01, M01, M02, and M03) for the observed (top) and randomized data. Blue lines represent the trend lines for each individual OTU, showing an overall negative bias in both observed and randomized data. F01 has the OTU shown in Panels A, B and C in larger red symbols.

**Supplemental Table 1: List of interacting OTUs.** A list of all significant interactions detected with time-lagged correlation including the greengenes OTU IDs of each interacting pair where OTU j (column A) impacts OTU i (column B), the p-value of the interaction (column C), whether the interaction was positive (facilitative) or negative (competitive) (column D), the taxonomic assignments of OTU i (column E) and OTU j (column F), the phylogenetic branch-length distance between OTU i and OTU j (column G), and the subject ID of the study participant in which the interaction was detected (column H).

